# Modeling the Degradation Effects of Autophagosome Tethering Compounds (ATTEC)

**DOI:** 10.1101/2020.04.01.021063

**Authors:** Hang Zhang, Ping An, Yiyan Fei, Boxun Lu

**Affiliations:** Department of Optical Science and Engineering, Shanghai Engineering Research Center of Ultra-Precision Optical Manufacturing, Key Laboratory of Micro and Nano Photonic Structures (Ministry of Education), Fudan University, Shanghai, China; State Key Laboratory of Medical Neurobiology and MOE Frontiers Center for Brain Science, School of Life Sciences, Fudan University, Shanghai, China

**Keywords:** Protein degradation, Autophagy, LC3, Bifunctional molecules, Drug discovery, Ternary complex, Simulation

## Abstract

Autophagy is a powerful protein degradation pathway with limited specificity. Our recent study proposed and demonstrated a potential strategy to harness autophagy to selectively degrade a specific pathogenic protein using autophagosome tethering compounds (ATTEC). ATTEC interact with both the target protein and the autophagosome protein LC3, and thus tether the target protein to the autophagosomes for subsequent degradation. The concentration-dependent curve of the target protein is U-shaped, but there has been lack of both kinetic and steady-state modeling of the degradation effects of ATTEC. Here we established a simplified model describing the kinetics and steady-state level of target protein, and characterized how compounds’ properties, especially binding affinities to LC3 and to the target protein, may influence their degradation effects.

## 1 Introduction

Selectively degradation of specific pathogenic proteins provides exciting strategies for drug discovery. Emerging new concepts such as PROTAC, ENDTAC, ATTEC are based on the design of “molecular glues” or bifunctional chimeric compounds that tether the target protein (protein of interest, POI) with a certain component of the protein degradation machinery (PDM)^[1-3]^. The formed trimer (POI·compound·PDM) then accelerates the POI degradation, leading its selective lowering.

Interestingly, the dose-dependent curves of the POI-compound relationship are U-shaped, with an optimal compound concentration for maximum lowering of POI and “hook” effects at higher compound concentrations^[1, 3, 4]^, different from traditional Boltzmann dose-dependent curves^[5]^. This is explained by the logic reasoning that when the compound concentration is too high, each compound molecule may interact with the POI and PDM separately, without tethering them together. Meanwhile, there has been a lack of mathematic modeling describing such effects. While modeling of the trimer formation has been published in a top tier journal^[6]^, it did not consider the degradation of the POI at all.

Such modeling has been challenging, because incorporating the degradation will greatly increase the calculation of the modeling. In addition, most of the existing degrader technologies are based on ubiquitination^[1]^, which is a complicated enzymatic reaction that is highly challenging to model^[7]^.

We have proposed and demonstrated a new degrader technology by harnessing autophagy for selective degradation using autophagosome tethering compounds (ATTEC)^[3]^. We demonstrated that compounds that interact with both the POI and the autophagosome protein LC3 may tether POI to autophagosomes for subsequent autophagic degradation^[3]^. Since ATTEC tether POI directly to the PDM without involvement of complicated enzymatic reactions, it may provide an ideal scenario for mathematic modeling of the degrader’s effects^[3]^. Here we describe a simplified model of the degradation effects of ATTEC, providing possible insights for understanding the dose-dependent data and potential clues for inventing better compounds.

## 2 Materials and methods

### 2.1 Modeling assumptions

Many parameters may influence the degradation kinetics and the dose-dependent effects^[8, 9]^, and considering all of them as variables may make the model highly complicated and difficult to resolve. Thus, we focused on the relationship between the degradation of the POI and the ATTEC’s affinity to the POI or LC3, and considered them as the only variables. Meanwhile, we estimated the values of all other parameters and considered them as constants to simplify the model as much as possible.

The basic principle of ATTEC is to enhance the turnover of POI by tethering the POI to the autophagosome, and thus we need to estimate the turnover rate (or half-life) of the POI and of the POI tethered to the autophagosome. When tethered to the autophagosome, the turnover rate of the POI could be estimated by the turnover rate of SQSTM1/p62, an autophagy receptor protein that is mainly degraded by autophagy^[10, 11]^ and has a half-life of about 6 hours^[12]^. Thus, we may roughly assume that the half-life of the POI tethered to the autophagosome is 6 hours. In our study demonstrating the concept of ATTEC, we observed increased degradation of mutant HTT protein (mHTT) induced by ATTEC. Thus, we used mHTT as the POI for our modeling, and its half-life has been reported to be around 33∼40 hours^[13-15]^. In order to facilitate the math calculation, we estimate the mHTT half-life as 36 hours, 6 times of the half-life of mHTT tethered to autophagosomes.

Besides the turnover rates, we also need to estimate the starting concentrations of POI (mHTT) and LC3 before compound treatment. It has been estimated that the cells have 2∼4 million (or a median of 3 million) protein molecules per cubic micron^[16]^. In addition, our previous proteomics data showed that the factions of total HTT (including both mHTT and wild-type HTT) and LC3 are 1/100,000 and 6/100,000 of total protein, respectively^[3]^. Based on the information above, we estimate that the starting concentrations of intracellular mHTT and LC3 are 33 and 400 nM, respectively.

In addition to the estimation of these biological parameters, we made some assumptions to further simplify the model. First, we assume that the LC3 level remains constant during the process of drug treatment. This is reasonable because experimental data showed that neither the LC3-I and the LC3-II levels were influenced by the ATTEC treatment^[3]^. In addition, it is consistent with the fact that LC3 is an abundant protein with fast turnover rate^[17, 18]^. We also assume that the total levels of ATTEC molecules remain constant. We recognize that this assumption is over-simplified. Meanwhile, it could be a reasonable assumption since the culture medium changed every day during the experiments, which may replenish the potential loss of the compound due to its degradation or metabolic changes. In addition, the experimental degradation rate of the compounds in free and in protein-bound forms are extremely difficult to determine and may add a great layer of complexity to our model. Thus, we focus on the target protein degradation only and assume a constant level of the compounds in cell culture.

Finally, to simplify the model, we assume that ATTEC’s binding to one protein does not influence its binding to the other, i.e., the LC3-bound ATTEC and free ATTEC have the same affinity to mHTT, and the mHTT-bound ATTEC and free ATTEC have the same affinity to LC3. We made this assumption to simply the model, which could be modified if additional evidence reveals a positive or negative coupling between the two proteins’ binding to ATTEC.

### 2.2 Kinetic equations and modeling

We considered the following chemical reactions for modeling,

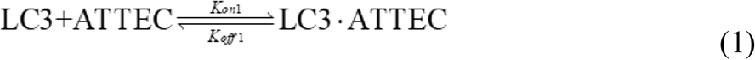

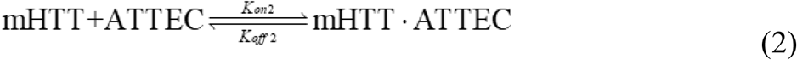

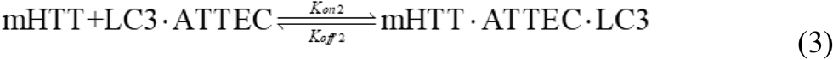

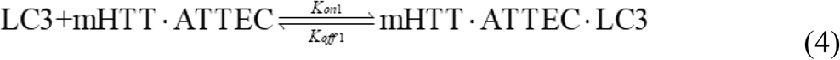

where the *K*_*on1*_ and *K*_*off1*_ are association rate constants and dissociation rate constants of the LC3 binding to the mHTT-bound ATTEC and free ATTEC, respectively; the *K*_*on2*_ and *K*_*off2*_ are the association rate constants and dissociation rate constants of the mHTT binding to LC3-bound ATTEC and free ATTEC, respectively.

Based on the chemical reactions above and the assumptions mentioned in the previous session, the kinetic equations were written as,

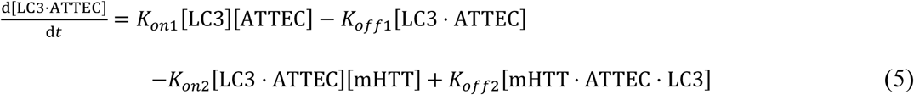

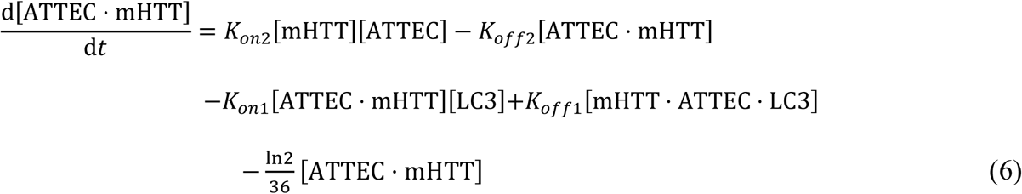

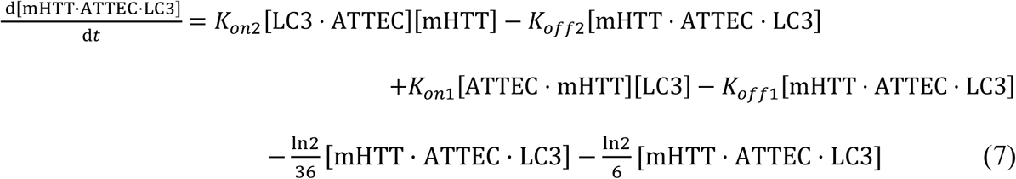

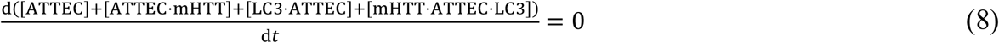

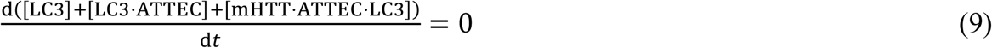

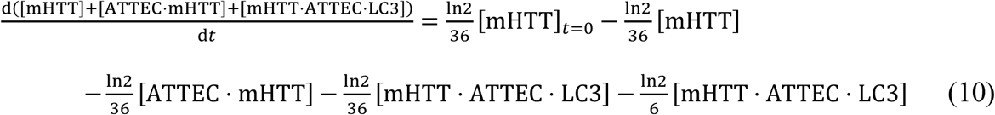

Eq. (5) is based on the kinetics of chemical reaction (1) and (3), whereas Eq. (6) is based on the kinetics of chemical reaction (2) and (4), as well as the degradation of the binary complex ATTEC·mHTT, which is assumed to be degraded at the same rate as free mHTT with a half-life of 36 hours. Note that the binary complexes participate in not only the reaction between free protein and free ATTEC (reaction (1) and (2)), but also the reaction to form ternary complexes (reaction (3) and (4)). Similarly, Eq. (7) is based on the kinetics of reaction (3) and (4), as well as the degradation of the ternary complex, which is subject to both the ATTEC-induced autophagic degradation with an estimated half-life of 6 hours and the endogenous degradation with an estimated half-life of 36 hours. The concentration of the ternary complex increases with binary complex association and decreases with ternary complex dissociation as well as its degradation. Eq. (8) and Eq. (9) are based on the assumption that the total concentration of LC3 and ATTEC remain constant over time. Eq. (10) is written based on the protein synthesis rate and degradation of several different mHTT-containing species. The mHTT is synthesized over time, and the synthesis rate is determined by the level of mutant *HTT* mRNA and the protein translation rate. These should not be changed by ATTEC, which only targets the mutant *HTT* protein but not mRNA to autophagic degradation. Thus, the protein synthesis rate should remain constant, which could be estimated by the fact that the mHTT remains at steady-state level before ATTEC treatment. Thus, based on the equilibrium of mHTT protein synthesis and degradation, the synthesis rate could be calculated by the mHTT half-life (36 hours) and its starting concentration ([mHTT]_*t*=0_) to be 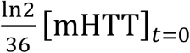. Similarly, the degradation of total mHTT by endogenous degradation system (ATTEC irrelevant) could be calculated based on a half-life of 36 hours, but the degradation rates are proportional to the real-time concentrations of each individual species. Thus, the ATTEC-irrelevant degradation rate of total mHTT equals 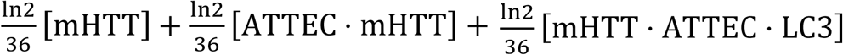. The ternary complex mHTT·ATTEC·LC3 is degraded additionally by the ATTEC-induced autophagic degradation, which is estimated to have a half-life of 6 hours and accounts for the 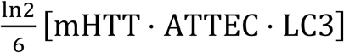 component in Eq. (10). Note that a minus sign is added to each degradation component to indicate a decrease of concentration over time.

### 2.3 Steady-state equations and modeling

In order to simulate and understand the dose-dependent curves of ATTEC-induced mHTT lowering, we need to calculate the total mHTT level at different compound concentrations based on steady-state equations shown as below:

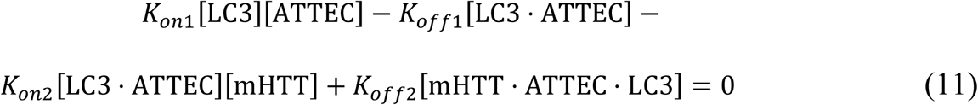

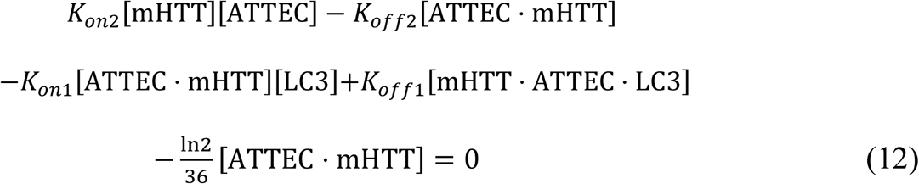

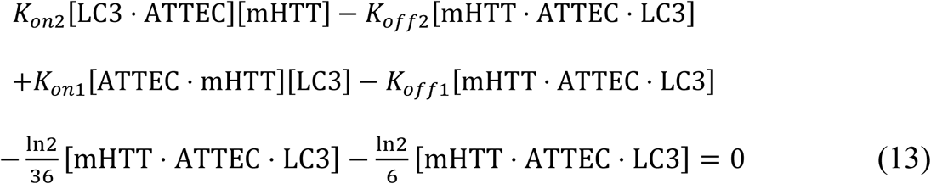

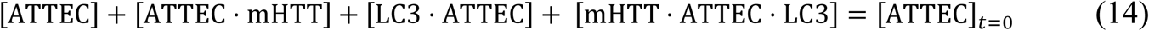

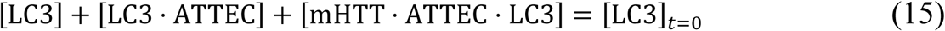

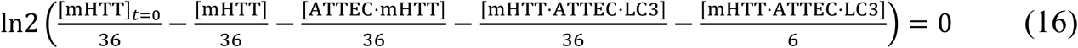

Eq. (11), (12) and (13) are based on the kinetic equations (5) through (7), whereas Eq. (14) and (15) are based on the assumption that the total ATTEC and total LC3 concentrations remain constant. For steady-state, the total mHTT reaches an equilibrium level, and this is described by Eq. (16).

## Results and discussion

### 3.1 Kinetic modeling of each specie involved in ATTEC

We first solved kinetic equations Eq. (5) to Eq. (10) with codes written with Matlab and simulated the kinetic changes of different species in the system. At the indicated kinetic parameters, the kinetic changes of each species were calculated and illustrated in Fig. 1. The kinetic parameters were estimated based on our experimental measurement of the GW5074 compound previously^[3]^.

**Fig. 1.**
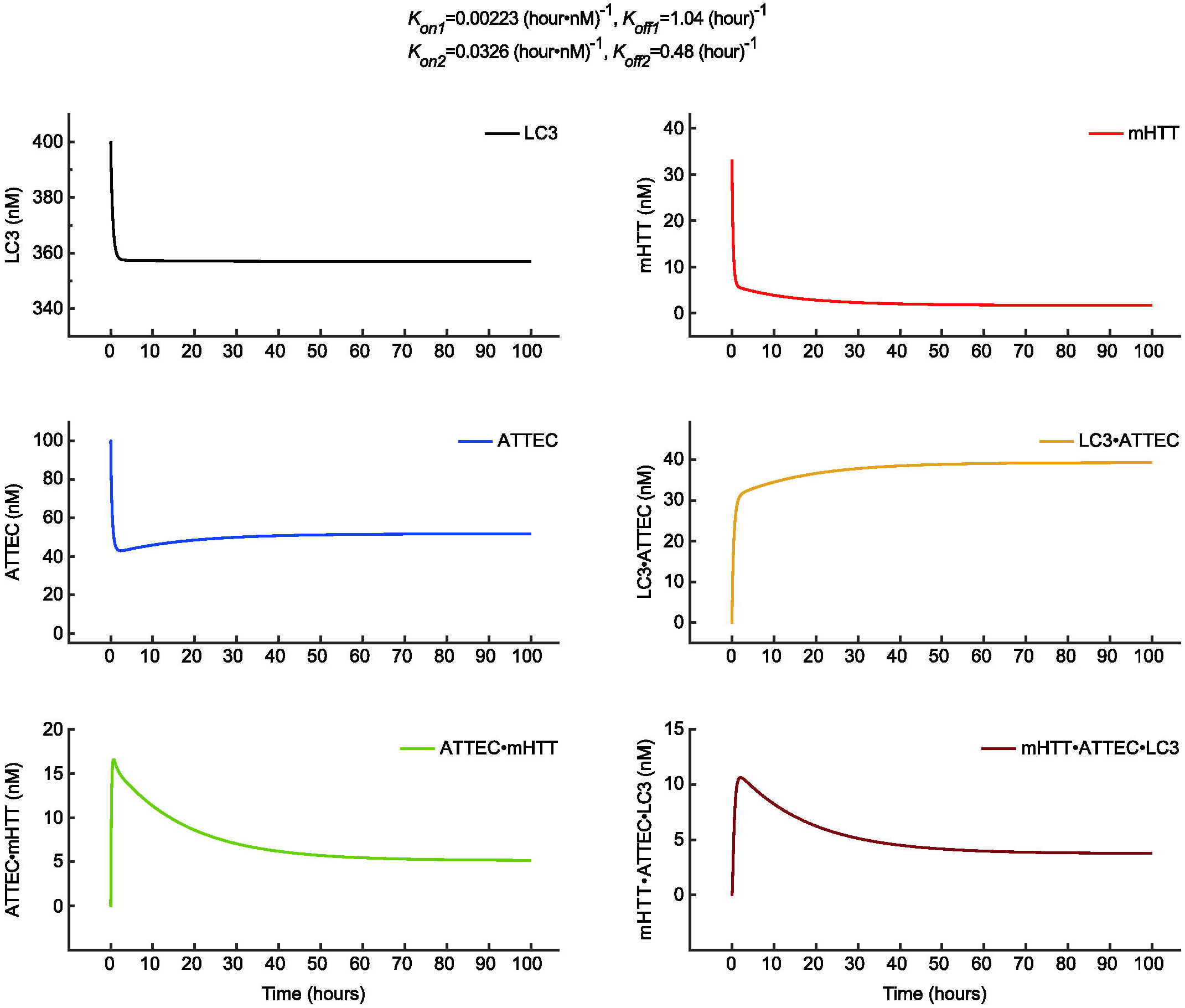
Kinetic modeling of the indicated species involved in ATTEC. The simulated concentrations of each of the indicated species were plotted over time. The species include the free LC3 protein (LC3), the free mHTT (mHTT), the free ATTEC molecule (ATTEC), the LC3·ATTEC binary complex (LC3·ATTEC), the ATTEC·mHTT binary complex (ATTEC·mHTT), and the mHTT·ATTEC·LC3 ternary complex (mHTT·ATTEC·LC3). The kinetics parameters used for modeling are presented on top of the figure. The ATTEC concentration for modeling is 100 nM.

Based on the kinetic simulation, the free mHTT (POI) and free LC3 protein had the fastest decay rates and reached steady-state within a few hours (Fig. 1, the mHTT panel and the LC3 panel), due to a fast formation of the ATTEC·mHTT and the ATTEC·LC3 binary complexes, respectively. The free ATTEC molecule had a rapid decay but bounced back a little at a much slower rate (Fig. 1, the ATTEC panel). The first decay phase is due to the fast formation of the ATTEC·LC3 and the ATTEC·mHTT binary complexes as well. The bounced back illustrates the recycle of ATTEC after degradation of the mHTT·ATTEC·LC3 proteins. The LC3·ATTEC complex had a rapid increase due to the formation of this binary complex (Fig. 1, the LC3·ATTEC panel). Both the ATTEC·mHTT binary complex and the mHTT·ATTEC·LC3 ternary complex increased rapidly and then decayed slowly to reach a steady-state level (Fig. 1, the ATTEC·mHTT panel and the mHTT·ATTEC·LC3 panel). The rapid increase phase is due to the formation of these complexes, and the slow decay phase of ATTEC·mHTT is due to the degradation of this complex as well as the consumption of this complex due to the formation of the ternary complex. The slow decay phase of the mHTT·ATTEC·LC3 complex is due to the degradation of this complex, and balanced by formation of this complex.

The major purpose of the model is to understand the kinetics of the total mHTT level, which is the sum of free mHTT, the binary ATTEC·mHTT complex, and the ternary mHTT·ATTEC·LC3 complex. Thus, we calculated the sum of the levels of these three species, and simulated the total mHTT kinetic curves (Fig. 2). The total mHTT level exhibited a decay over time, reaching a steady-state after ∼30 to ∼150 hours depending on the kinetic parameters (Fig. 2). We then investigated how the kinetic parameters may influence the kinetic curve of the total mHTT level. The *K*_*on1*_ and *K*_*off1*_ values determine the kinetics of the formation of ATTEC·LC3 complex and mHTT·ATTEC·LC3 complex. Larger *K*_*on1*_ or smaller *K*_*off1*_, will lead to faster ATTEC·LC3 and mHTT·ATTEC·LC3 formation. Consistent with this, the total mHTT level decays faster and reaches a lower steady-state level with larger *K*_*on1*_ and smaller *K*_*off1*_ (Fig. 2). Noticeably, the curves are more sensitive to *K*_*on1*_ changes in a lower value range (between 0.000223 and 0.00223 (hour·nM)^-1^) but to *K*_*off1*_ changes in a higher value range (between 10.4 and 1.04 (hour)^-1^). Similar observations were made on the influences by *K*_*on2*_, *K*_*off2*_, which determines the kinetics of the formation of ATTEC·mHTT and mHTT·ATTEC·LC3 complexes (Fig. 2). Under all conditions, the total mHTT level at 48 hours is very close to the steady-state level, suggesting that the 48 hours compound treatment time used in the previously published experiments is reasonable^[3]^.

**Fig. 2.**
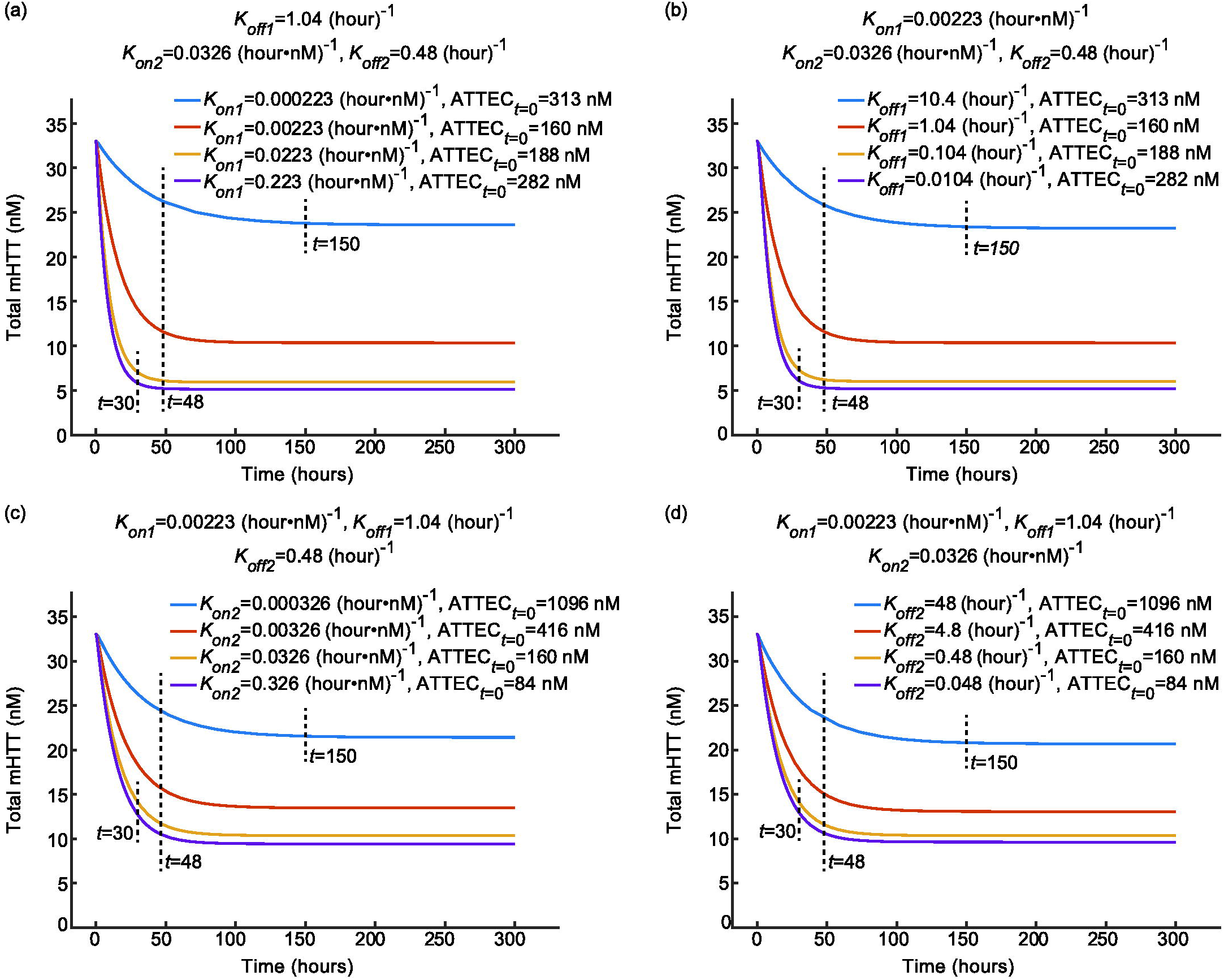
Kinetic modeling of the total mHTT concentrations. **(a)** The simulated total mHTT concentration-time relationship with the indicated set of *K*_*on1*_ values when *K*_*off1*_, *K*_*on2*_, and *K*_*off2*_ are fixed. **(b)** Similar to (a), but with the indicated set of *K*_*off1*_ values when fixing all other kinetic parameters. **(c)** Similar to (a), but with the indicated set of *K*_*on2*_ values when fixing all other kinetic parameters. **(d)** Similar to (a), but with the indicated set of *K*_*off2*_ values when fixing all other kinetic parameters. Note that the ATTEC concentrations used for each simulation are the optimal concentrations to achieve highest D_max_ under each condition (see Fig. 4). The dashed lines indicate the time needed to reach steady-state levels.

### 3.2 Modeling of the dose-dependent curves for ATTEC

We then solved the steady-state equations and simulated the dose-dependent curves of ATTEC. All the ATTEC curves were U-shaped (Fig. 3), consistent with the autophagosome-tethering mechanism: a sufficient concentration of ATTEC is needed, whereas excessively high compound concentrations may lead to binding of ATTEC to LC3 and mHTT separately, without tethering the two together. Since at the same *K*_*d*_ (the equilibrium dissociation constant) values, the dose-dependent curves change very little with *K*_*on*_ and *K*_*off*_ varying two orders of magnitude (Fig. 3), we focused on analyzing *K*_*d*_ values’ effect on ATTEC’s degradation efficiency.

**Fig. 3.**
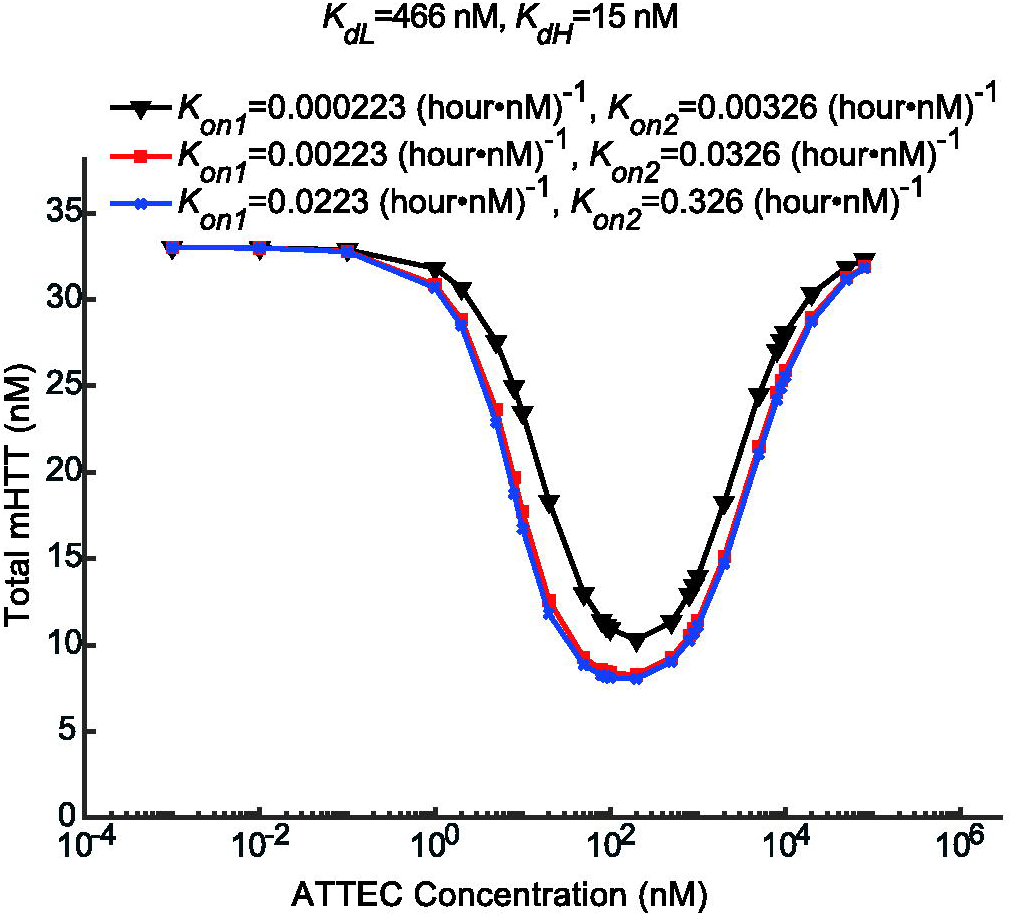
Modeling of the dose-dependent curves for ATTEC with fixed *K*_*d*_ values. The simulated relationship between the steady-state concentrations of total mHTT and the concentrations of ATTEC molecules with fixed *K*_*dH*_ and *K*_*dL*_ by varying *K*_*on1*_, *K*_*off1*_, *K*_*on2*_, and *K*_*off2*_ accordingly.

We simulated the dose-dependent curves with different sets of compound-protein affinity parameters (Fig. 4, *K*_*dH*_ for ATTEC’s affinity to mHTT, and *K*_*dL*_ for ATTEC’s affinity to LC3). The shape of dose-dependent curve highly resembles experimental data that we published previously^[3]^. The target protein mHTT could only be lowered to a certain degree, the extend of lowering is referred to as maximum degradation (D_max_). D_max_ is reached only at optimal concentrations of ATTEC. It is clear that both D_max_ and optimal concentrations for degradation are influenced by the *K*_*d*_ values (Fig. 4), which are further analyzed as below.

**Fig. 4.**
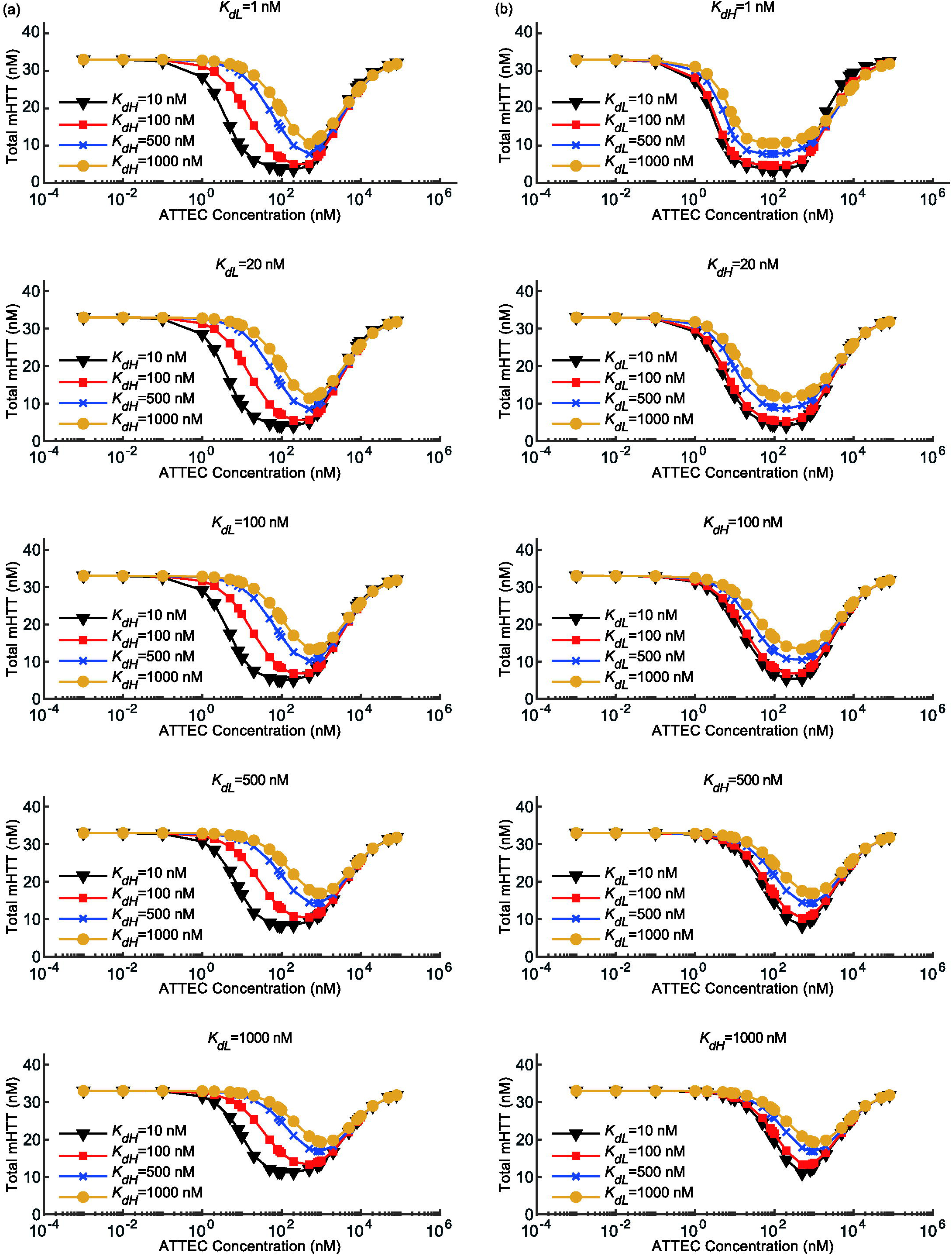
Modeling of the dose-dependent curves for ATTEC. The relationship between the steady-state concentrations of total mHTT and the concentrations of ATTEC molecules were simulated. **(a)** The ATTEC dose-dependent curves with a set of *K*_*dH*_ and a fixed *K*_*dL*_ in each panel. **(b)** similar to (a), but with a set of *K*_*dL*_ and a fixed *K*_*dH*_ in each panel.

### 3.3 Relationship between affinity (*K*_*d*_) and maximum degradation (D_max_)

In order to visualize the relationship between D_max_ and *K*_*d*_ values, we plotted the D_max_-*K*_*dH*_ relationship assuming several different *K*_*dL*_ values (Fig. 5a). The *K*_*dH*_ increase leads to decreased D_max_ values, consistent with the prediction that lowering affinity (higher *K*_*d*_) to the POI (mHTT) leads to less efficient degradation. Meanwhile, the curves are relatively flat within the range of 1∼100 nM *K*_*dH*_, and drop rapidly within the range of 100∼10000 nM *K*_*dH*_ (Fig. 5a), suggesting that the D_max_ is sensitive to *K*_*dH*_ only in the low affinity range (high *K*_*dH*_ values at 100 ∼ 10000 nM). The D_max_-*K*_*dL*_ relationship showed highly similar results (Fig. 5b), suggesting that the D_max_ is sensitive to *K*_*dL*_ only in the low affinity range as well. Thus, while relatively high affinity of ATTEC to POI (mHTT) and LC3 is desired to achieve higher D_max_, it may not be necessary to make huge efforts to achieve extremely high affinity (such as ∼1 nM *K*_*d*_), because *K*_*d*_ values within 1∼100 nM lead to similar D_max_.

**Fig. 5.**
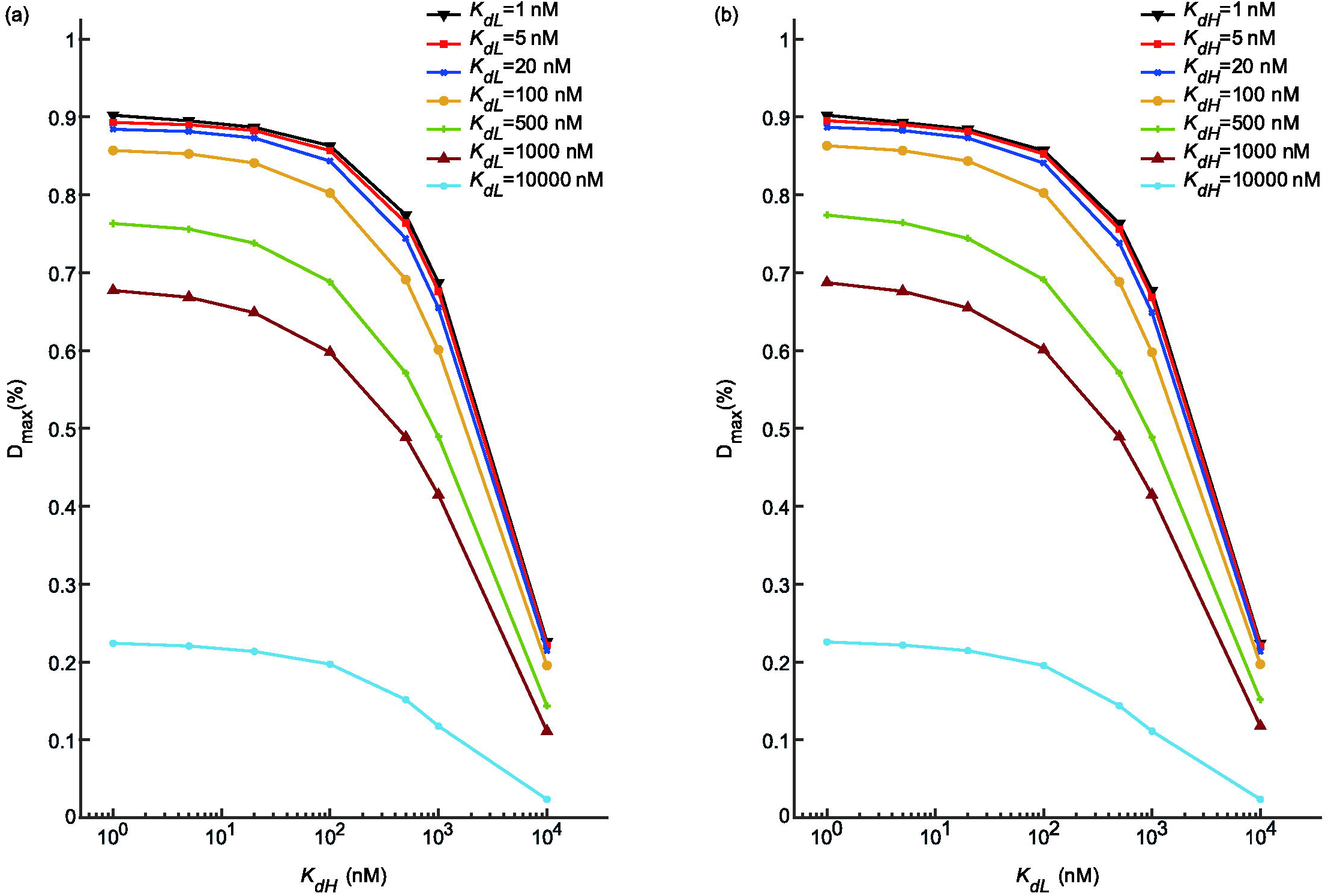
The maximum degradation D_max_ changes with various of affinity *K*_*dL*_ and *K*_*dH*_. **(a)** The simulated Dmax-*K*_*dH*_ relationship assuming the indicated *K*_*dL*_ values. **(b)** The simulated Dmax-*K*_*dL*_ relationship assuming the indicated *K*_*dH*_ values.

### 3.4 Relationship between affinity (*K*_*d*_) and effective compound concentrations

As mentioned above, the dose-dependent curve of ATTEC is unlikely the common Boltzmann curves which fit most chemical responses. Instead, the ATTEC dose-dependent curve is U-shaped, and an optimal concentration range is desired for efficient degradation. This optimal range could be influenced by ATTEC’s affinity to both POI (mHTT) and LC3, as illustrated by our modeling data assuming several different sets of *K*_*d*_ values (Fig. 4). Resolving the relationship between optimal concentration range and the *K*_*d*_ values may guide compound optimization.

In order to visualize this relationship, we plotted three different ATTEC concentrations against the *K*_*d*_ values. ATTEC-D_max_ indicates the ATTEC concentration needed to reach the maximum degradation effects. ATTEC-D_left_ and ATTEC-D_right_ indicate the lower and higher concentrations needed to reach half of the maximum degradation effect, respectively.

We first plotted these concentrations against *K*_*dH*_, assuming several different *K*_*dL*_ values (Fig. 6a). The general trend is that ATTEC-D_max_, ATTEC-D_left_ and ATTEC-D_right_ all increases with *K*_*dH*_ (Fig. 6a), suggesting that higher ATTEC concentrations are needed to achieve efficient degradation when ATTEC’s affinity to mHTT decreases, represented by increased *K*_*dH*_ values. In addition, the size of the effective concentration window (Fig. 6a, distance between the yellow and red curves, log scale) also decreases as *K*_*dH*_ increases, especially at larger *K*_*dH*_ range, suggesting that the effective concentration range is larger when ATTEC’s affinity to POI (mHTT) is high. Finally, the ATTEC-D_left_ curves generally have a larger slope, suggesting that the lower concentration required to achieve half of the maximum degradation effect is more sensitive to ATTEC’s affinity to POI (mHTT).

**Fig. 6.**
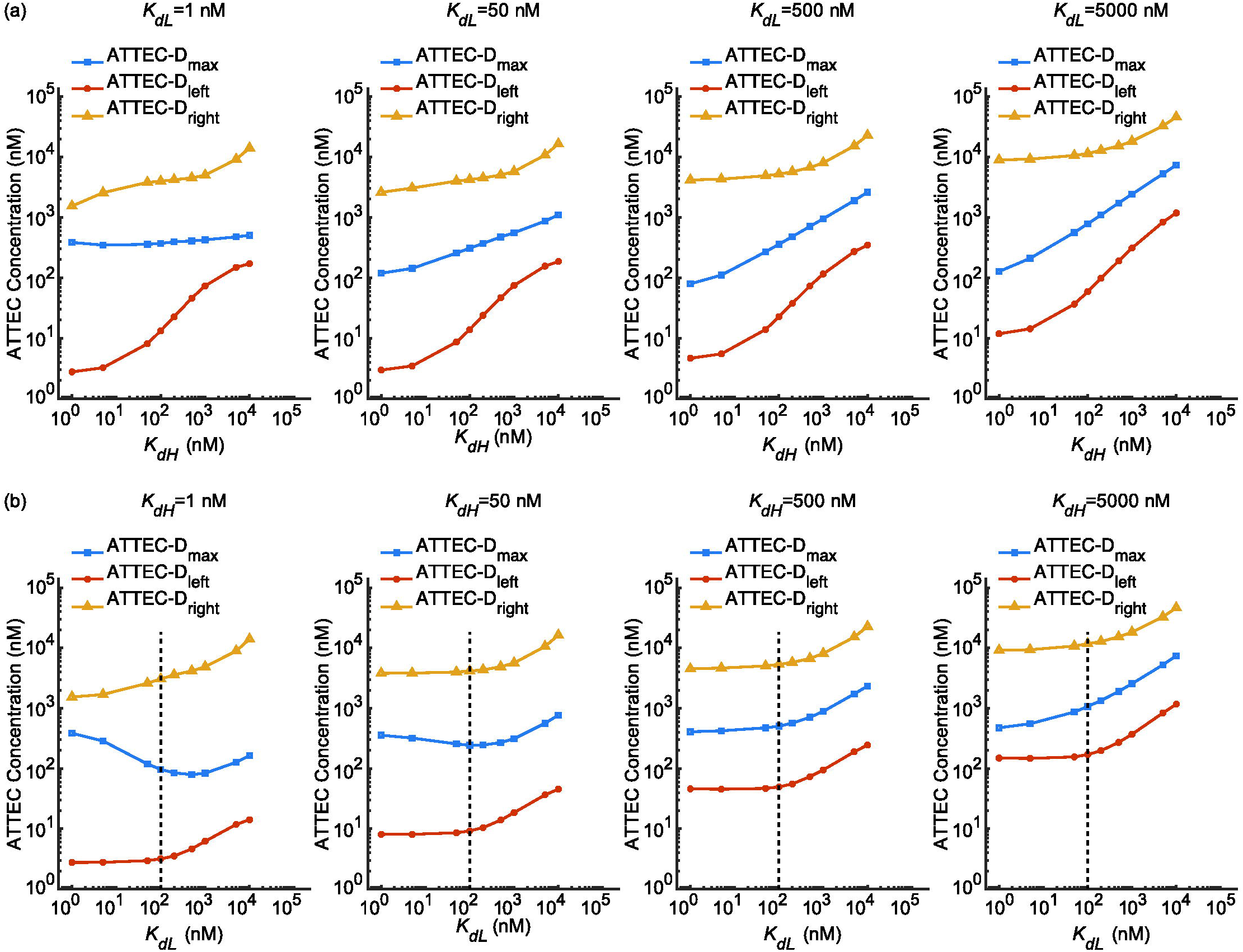
The effective concentration ranges of ATTEC for effective degradation. The concentrations needed to achieve maximum degradation ATTEC-D_max_ or half-maximum degradation ATTEC-D_left_ and ATTEC-D_right_ at different sets of *K*_*dL*_ and *K*_*dH*_ values are plotted. **(a)** The ATTEC-D_max_, ATTEC-D_left_, and ATTEC-D_right_ concentrations assuming different *K*_*dH*_ at a fixed *K*_*dL*_ in each panel. **(b)** similar to (a), but assuming different *K*_*dL*_ at a fixed *K*_*dH*_ in each panel. The dashed lines indicate 100 nM, below which *K*_*dL*_ has little influence on the effective concentration range.

We then plotted ATTEC-D_max_, ATTEC-D_left_ and ATTEC-D_right_ against *K*_*dL*_. Interestingly, the curves looked different from the ones plotted against *K*_*dH*_. The major difference is that the size of the effective concentration window (Fig. 6b, distance between the yellow and red curves, log scale) is insensitive to *K*_*dL*_ values, suggesting that the ATTEC’s affinity to LC3 is not a major factor contributing to the size of effective concentration range. Second, the ATTEC-D_max_ curve (Fig. 6b, blue) is not always monotonic increasing, and optimal *K*_*dL*_ between 100∼1000 nM is required to reach lowest ATTEC-D_max_, at least when *K*_*dH*_ is low (Fig. 6b, *K*_*dH*_ = 1 nM or 50 nM). Thus, for an attempt to reach maximum degradation effects with low ATTEC concentrations, it seems that high affinity of ATTEC to LC3 is not always desired. Finally, for *K*_*dL*_ below 100 nM (Fig. 6b, left to the dashed lines in each panel), the ATTEC-D_left_ and ATTEC-D_right_ were largely unaffected, whereas they did increase when *K*_*dL*_ is larger than 100 nM. Thus, in order to induce efficient degradation at lower ATTEC concentrations, a higher affinity to LC3 (lower *K*_*dL*_) is desired only when *K*_*dL*_ is larger than 100 nM. Further increasing the affinity to LC3 may not be necessary, at least in conditions assumed in our modeling.

## 4 Conclusions and discussion

In this study we presented the first simplified model of ATTEC-induced lowering of POI. While the model is over-simplified, the shape of dose-dependent curves largely fit with experimental data^[3]^. We further studied the influence of ATTEC’s affinities to POI (mHTT) and LC3, and observed some interesting properties. In general, the D_max_ is negatively correlated with both *K*_*dH*_ and *K*_*dL*_, but not so sensitive when the *K*_*d*_ values are below 100 nM (Fig. 5). Lower *K*_*dH*_ values are also required to achieve desired effective concentration range (Fig. 6a), but lower *K*_*dL*_ values may not be necessary, especially when *K*_*dL*_ is smaller than 100 nM (Fig. 6b). Thus, optimizing the compounds to reach a higher affinity to POI (mHTT) is more important, and an attempt to reach extremely high affinity to LC3 (<100 nM) may not be necessary. These predictions are based on many assumptions including the estimation of the starting concentrations of mHTT and LC3 proteins, and certainly need further confirmation by experiments. Meanwhile, as the first modeling for ATTEC-induced degradation, and even for ternary complex-induced degradation, it may still provide many insights into the properties of such degradation and possible factors to consider for compound optimization.

## Conflict of interest

The authors declare that they have no conflict of interest.

## Acknowledgement

This work was supported by National Natural Science Foundation of China: 81925012, National Natural Science Foundation of China: 81870990, National Natural Science Foundation of China: 31961130379 and Newton Advanced Fellowship: NAF_R1_191045.

## References

[1] Bondeson DP, Mares A, Smith IE, et al. Catalytic in vivo protein knockdown by small-molecule protacs. Nature chemical biology, 2015, 11: 611–617

[2] Nalawansha DA, Paiva SL, Rafizadeh DN, et al. Targeted protein internalization and degradation by endosome targeting chimeras (endtacs). ACS central science, 2019, 5: 1079–1084

[3] Li Z, Wang C, Wang Z, et al. Allele-selective lowering of mutant htt protein by htt-lc3 linker compounds. Nature, 2019, 575: 203–209

[4] Winter GE, Buckley DL, Paulk J, et al. Drug development. Phthalimide conjugation as a strategy for in vivo target protein degradation. Science (New York, NY), 2015, 348: 1376–1381

[5] Dubois JM, Ouanounou G, Rouzaire-Dubois B. The boltzmann equation in molecular biology. Progress in biophysics and molecular biology, 2009, 99: 87–93

[6] Douglass EF, Jr., Miller CJ, Sparer G, et al. A comprehensive mathematical model for three-body binding equilibria. Journal of the American Chemical Society, 2013, 135: 6092–6099

[7] Schapira M, Calabrese MF, Bullock AN, et al. Targeted protein degradation: Expanding the toolbox. Nature reviews Drug discovery, 2019, 18: 949–963

[8] Zhang Y, Loh C, Chen J, et al. Targeted protein degradation mechanisms. Drug discovery today Technologies, 2019, 31: 53–60

[9] Crisp MJ, Mawuenyega KG, Patterson BW, et al. In vivo kinetic approach reveals slow sod1 turnover in the cns. The Journal of clinical investigation, 2015, 125: 2772–2780

[10] Johansen T, Lamark T. Selective autophagy mediated by autophagic adapter proteins. Autophagy, 2011, 7: 279–296

[11] Myeku N, Figueiredo-Pereira ME. Dynamics of the degradation of ubiquitinated proteins by proteasomes and autophagy: Association with sequestosome 1/p62. The Journal of biological chemistry, 2011, 286: 22426–22440

[12] Bjorkoy G, Lamark T, Brech A, et al. P62/sqstm1 forms protein aggregates degraded by autophagy and has a protective effect on huntingtin-induced cell death. The Journal of cell biology, 2005, 171: 603–614

[13] Tsvetkov AS, Arrasate M, Barmada S, et al. Proteostasis of polyglutamine varies among neurons and predicts neurodegeneration. Nature chemical biology, 2013, 9: 586–592

[14] Wu P, Lu MX, Cui XT, et al. A high-throughput-compatible assay to measure the degradation of endogenous huntingtin proteins. Acta pharmacologica Sinica, 2016, 37: 1307–1314

[15] Fu Y, Wu P, Pan Y, et al. A toxic mutant huntingtin species is resistant to selective autophagy. Nature chemical biology, 2017, 13: 1152–1154

[16] Milo R. What is the total number of protein molecules per cell volume? A call to rethink some published values. BioEssays : news and reviews in molecular, cellular and developmental biology, 2013, 35: 1050–1055

[17] Ni HM, Bockus A, Wozniak AL, et al. Dissecting the dynamic turnover of gfp-lc3 in the autolysosome. Autophagy, 2011, 7: 188–204

[18] Moulis M, Vindis C. Methods for measuring autophagy in mice. Cells, 2017, 6:

